# Bee and floral traits affect the characteristics of the vibrations experienced by flowers during buzz-pollination

**DOI:** 10.1101/494690

**Authors:** Blanca Arroyo-Correa, Ceit Beattie, Mario Vallejo-Marín

**Affiliations:** Biological and Environmental Sciences, School of Natural Sciences, University of Stirling. Stirling, Scotland. FK9 4LA.; Integrative Ecology Group, Estación Biológica de Doñana (EBD-CSIC), Sevilla, Spain

## Abstract

During buzz pollination, bees use their indirect flight muscles to produce vibrations that are transmitted to the flowers and result in pollen release. Although buzz pollination has been known for >100 years, we are still in the early stages of understanding how bee and floral characteristics affect the production and transmission of floral vibrations. Here we analysed floral vibrations produced by four closely related bumblebee taxa (*Bombus spp.*) on two buzz-pollinated plants species (*Solanum spp.*). We measured floral vibrations transmitted to the flower to establish the extent to which the mechanical properties of floral vibrations depend on bee and plant characteristics. By comparing four bee taxa visiting the same plant species, we found that peak acceleration (PA), root mean-squared acceleration (RMS) and frequency varies between bee taxa, but that neither bee size (intertegular distance) or flower biomass (dry weight) affect PA, RMS or frequency. A comparison of floral vibrations of two bee taxa visiting flowers of two plant species, showed that, while bee species affects PA, RMS and frequency, plant species affects acceleration (PA and RMS) but not frequency. When accounting for differences in the transmission of vibrations across the two types of flowers, using a species-specific “coupling factor”, we found that RMS acceleration and peak displacement does not differ between plant species. This suggests that bees produce the same initial acceleration in different plants but that transmission of these vibrations through the flower is affected by floral characteristics.

**Summary statement:** We show that buzz-pollinating bumblebees differ in the type of vibrations produced while visiting the same flower, and that floral species affects the transmission properties of those vibrations.

## Introduction

Buzz pollination is a fascinating interaction between flowers with specialised morphologies and bees that use vibrations to remove pollen (Buchmann, 1983; Vallejo-Marin, in review). Buzz pollination is by no means rare. The ability to produce vibrations to remove pollen from flowers has evolved at least 45 times in the evolutionary history of bees, and it is estimated that approximately 6% of flowering plants are buzz-pollinated (Cardinal et al., 2018). In most buzz-pollinated plants, pollen is kept locked inside tubular, non-dehiscent anthers (i.e., poricidal anthers; Buchmann, 1983; Harris, 1905), or inside closed corolla tubes with small apical openings (Corbet and Huang, 2014; Kawai and Kudo, 2008; Macior, 1968) that restrict direct access to pollen. Pollen grains in buzz-pollinated flowers are most efficiently removed by bees that are capable of using floral vibrations—also called “sonications” or “buzzes”—to release pollen from poricidal anthers and flowers (De Luca and Vallejo-Marín, 2013). Despite the widespread occurrence of the ability to produce floral vibrations among bees (Hymenoptera: Apoidea: Anthophila), little is known about why some bee species buzz-pollinate (e.g., carpenter bees, bumblebees, several sweat bees), while others seem incapable of doing so (e.g., honeybees and most leaf-cutter bees) (Cardinal et al., 2018; De Luca and Vallejo-Marín, 2013). Moreover, we are still in the early stages of understanding the extent to which floral vibrations produced by bees vary between bee species and between different types of plants.

The production of floral vibrations by bees is a relatively stereotyped behaviour (De Luca and Vallejo-Marín, 2013). During a typical buzz-pollinating visit, a bee embraces one or more poricidal anthers, holding to the anthers using its mandibles, curls its body around the anthers, and produces one or more bursts of vibrations that can be heard by a human observer as “buzzes” of a higher pitch than that buzz heard during flight (Macior, 1964; Russell et al., 2016) (Fig. 1). A bee generates floral vibrations using its thoracic musculature, which causes rapid deformation of the thorax while the wings remain undeployed (King and Buchmann, 2003). These vibrations are transmitted as substrate-borne vibrations to the anthers through the mandibles/head, thorax and abdomen of the bee (King, 1993; Vallejo-Marin, in review). The vibrations result in pollen ejection from the anther tips (Fig. 1), probably as a consequence of energy transfer from the bee to the pollen grains (Buchmann and Hurley, 1978), which should be a function of the mechanical properties of the substrate-borne vibrations as well as the characteristics of both bee and flower (Vallejo-Marin, in review). Determining exactly what characteristics of floral vibrations are most important for pollen release is an area that requires further theoretical and empirical work.

**Figure 1.**
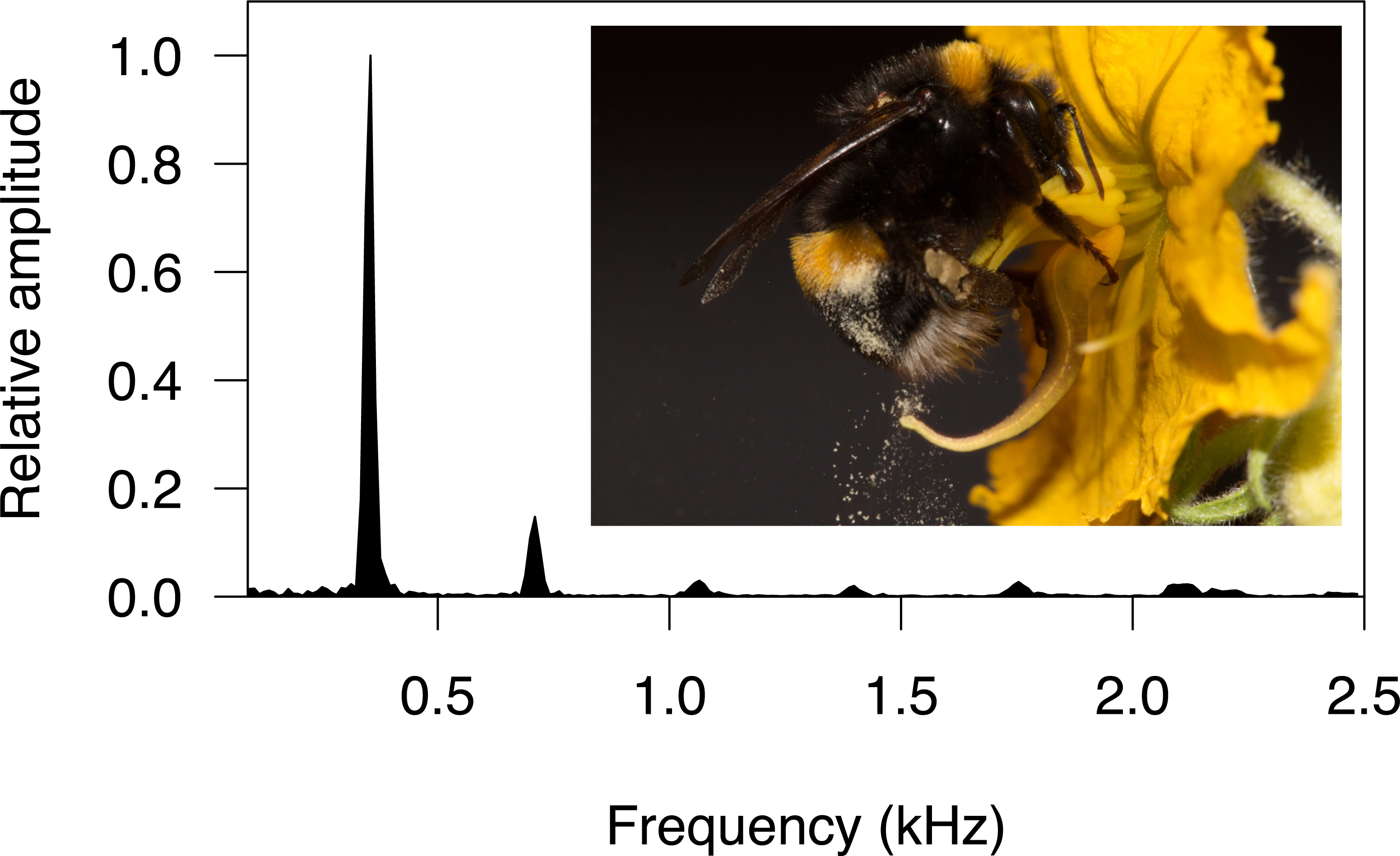
Frequency spectrum of a single floral vibration (shown in Fig. 2B) of the buff-tailed bumblebee (*Bombus audax*), on a flower of the buzz-pollinated herb buffalo bur (*Solanum rostratum*). The peak with the highest relative amplitude is the dominant frequency, which often corresponds to the fundamental frequency of the vibration (355 Hz in this example). The inset shows how the floral vibrations produced by the bumblebee result in pollen ejection from pores located at the tipof the anthers.

Floral vibrations produced by bees are a type of substrate-borne vibration and can be described by biophysical properties such as duration, frequency and amplitude (Cocroft and Rodríguez, 2005; Mortimer, 2017). During a single visit, a bee may produce one to several “buzzes”, but each of them can be itself made from very short vibrations (tens to hundreds of milliseconds) (Vallejo-Marin, in review) (Fig. 2a), although a single vibration may go uninterrupted for longer (seconds) (Buchmann and Cane, 1989). Floral vibrations contain multiple harmonics (Buchmann et al., 1977), although usually the fundamental frequency (first harmonic) contributes most to the overall magnitude of substrate-borne floral vibrations (De Luca and Vallejo-Marín, 2013; King, 1993). In such cases, the fundamental frequency is also the dominant or peak frequency (Fig. 1). The fundamental frequency of floral vibrations typically ranges between 100 and 500 Hz (De Luca and Vallejo-Marín, 2013). Floral vibrations can also be characterised by their amplitude (Figs. 2b and 2c). Amplitude describes the magnitude of oscillatory movement, and can itself be defined in terms of acceleration, velocity and displacement (Sueur, 2018). In many types of oscillatory movement, frequency, acceleration, velocity and amplitude are interrelated, and a full description of an oscillation thus requires knowledge of the absolute value of more than one of these variables. However, assuming simple harmonic oscillations it is possible to use two of these variables (e.g., frequency and acceleration) to calculate another one (e.g., displacement) (Vallejo-Marin, in review). An approach often taken to characterise floral vibrations during buzz pollination is to use acoustic recordings. Audio recordings provide accurate estimates of some aspects of substrate-borne vibrations, such as frequency and duration, but are less reliable in estimating the amplitude component (De Luca et al., 2018). As floral vibrations are in essence a substrate-borne phenomenon, measurements of the vibrations experienced by flowers thus require the vibrations transmitted to the flowers to be assessed directly using, for example, accelerometers or laser vibrometers (Vallejo-Marin, in review). To date, very few studies have attempted to compare floral vibrations produced by different species of bee visiting different species of plant using direct measurements of substrate-borne vibrations.

**Figure 2.**
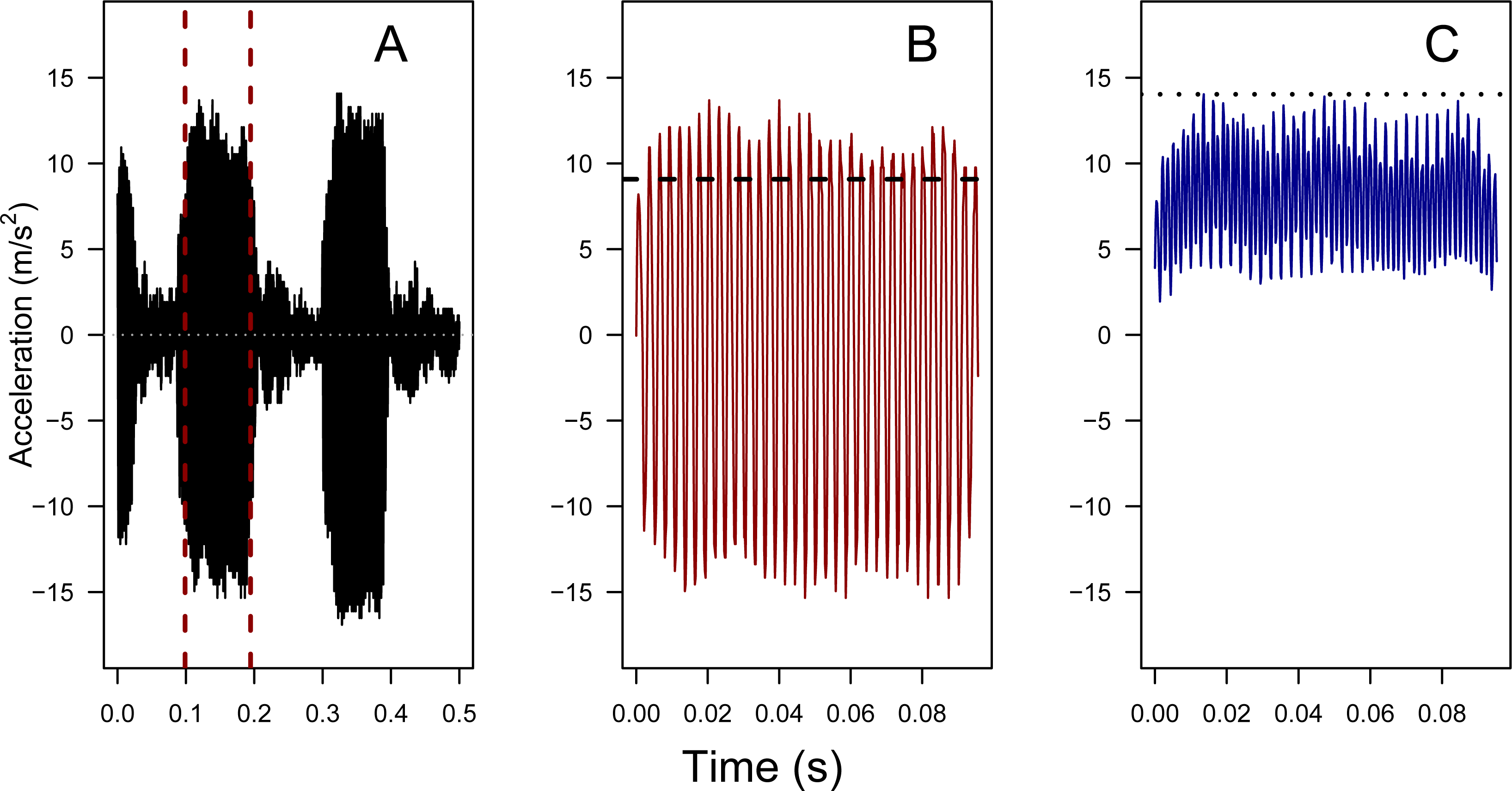
Floral vibration of Bombus terrestris ssp. audax on a flower of Solanum rostratum. **(A)** Oscillogram of the recorded floral vibration. The vertical dashed lines indicate the region of the vibration selected for analysis. **(B)** Zoom-in of the selected vibration. Root Mean Square acceleration (RMS) is calculated from these data and shown with a horizontal dashed line. **(C)** Amplitude envelope of the vibration shown in (B), calculated with a window size of two and a 75% overlap. Maximum peak acceleration was calculated from this amplitude envelope and is shown with a dashed line.

Our study focuses on bumblebees (*Bombus spp.* Latreille), which are well-known for their ability to buzz-pollinate and are important buzz-pollinators in both temperate and tropical regions (De Luca et al., 2014; Mesquita-Neto et al., 2018; Rosi-Denadai et al., 2018). There are approximately 250 species of bumblebee (*Bombus*: Bombini) worldwide, comprising bees of medium to very large size (9-22mm long) (Michener, 2000). Besides their important role as pollinators in natural systems, some species of bumblebees are commercially bred and used widely around the world for the pollination of some crops, including soft fruits such as raspberries (Lye et al., 2011), and buzz-pollinated plants such as tomatoes (Morandin et al., 2001). Previous work has shown that bumblebee species differ in the acoustic frequency of floral vibrations produced while visiting buzz-pollinated flowers (Corbet and Huang, 2014; De Luca et al., submitted; De Luca et al., 2014; Switzer and Combes, 2017), but it remains unclear how other characteristics of substrate-borne floral vibrations (e.g., acceleration and displacement) vary between bumblebee species. De Luca et al. (2013) found a positive correlation between body size and amplitude (peak velocity amplitude) of substrate-borne floral vibrations within a single bumblebee species. Frequency of floral vibrations (estimated from acoustic recordings) usually shows no correlation with body size within bumblebee species visiting a single plant (reviewed in De Luca et al., submitted), and show conflicting patterns when comparing the same bumblebee species visiting different plants (Corbet and Huang, 2014; Switzer and Combes, 2017). Although wingbeat frequency during flight is negatively associated with body size, the relationship between body size and the acoustic component of floral vibrations seems to be much weaker (De Luca et al., submitted). Comparisons of the properties of substrate-borne vibrations produced by different species of bee in the same flower and among different flowers are needed to better understand how bee species and individual characteristics influence the mechanical vibrations transmitted during buzz pollination.

One of the largest gaps in our current knowledge on buzz pollination is the effect of plant characteristics on the transmission of bee vibrations. Characteristics of the flower such as mass, stiffness, geometry and other material properties of anthers and associated floral structures are expected to affect the transmission of vibrations (Michelsen et al., 1982). Studies of animal communication using plant-borne substrates have shown that plant architecture (e.g., branch diameter, number of internodes between caller and receiver) affect the transmission of insect vibrations (Gibson and Cocroft, 2018). However, it is less clear whether plant characteristics influence the vibrations transmitted over short distances within a single flower. To date, only one study has quantitatively compared the transmission of floral vibrations among different plant species. King (1993) studied the differences in the amplitude of floral vibrations, measured as peak acceleration amplitude (PA) produced by bumblebees visiting flowers of *Symphytum sp.* (comfrey) and *Actinidia deliciosa* (kiwi). King calculated the ratio between the PA of vibrations of known amplitude applied to the anthers using a shaker table, with the PA of vibrations detected in the stem adjacent to the flower measured with an accelerometer. The ratio, which he called “coupling factor”, provides an estimate of the attenuation observed between the source of the vibration and the accelerometer, and is expected to be a function of the filtering and attenuating properties of the flower and intervening plant tissue (Cocroft and Rodríguez, 2005; Mortimer, 2017). King showed that the coupling factor differs between species with radically distinct floral morphologies (comfrey = 26.4; kiwi = 182). The results from this pioneering study raise two simple, but key points. First, plant identity and floral characteristics mediate the transmission of vibrations through the flower. Second, signal attenuation must be taken into account when trying to infer the characteristics of vibrations experienced by the anthers from measurements taken in other parts of the flower (e.g., stem, pedicel, calyx, petals).

Here, we investigate the characteristics of substrate-borne floral vibrations produced by four closely related bumblebee taxa (*Bombus s.s.*) (Cameron et al., 2007) on two buzz-pollinated plants in *Solanum* L. Section *Androceras* (Solanaceae). We used an accelerometer to collect measurements of acceleration and frequency from three taxa in the *Bombus terrestris* species aggregate (*B. terrestris, B. terrestris ssp. audax*, and *B. terrestris ssp. canariensis*) as well as in *B. ignitus,* while visiting flowers of *Solanum rostratum* and *S. citrullifolium*. We used these data to address four questions: (1) Do the properties of substrate-borne floral vibrations differ between bumblebee taxa when visiting flowers of the same plant species? (2) Do floral vibrations produced by the same bee taxon depend on the species of flower visited? (3) Do closely related plant species differ in their transmission properties of substrate-borne vibrations, as estimated using King’s coupling factor? And finally, (4) does bee size and/or floral biomass influence the characteristics of floral-borne vibrations transmitted to the flower during buzz pollination?

## Materials and Methods

### Bee material

We studied four closely related bumblebee taxa in *Bombus* subgenus *Bombus* Latreille (Hymenoptera, Apidae). All bee colonies used were reared and supplied by Biobest (Westerlo, Belgium). Three taxa belong to the *B. terrestris* L. species aggregate. Two of them are subspecies *B. terrestris ssp. audax* Harris (hereafter *B. audax*) and *B. terrestris ssp. canariensis* Pérez (hereafter B. *canariensis*) (Rasmont et al., 2008). These two taxa have different native distributions, with *B. audax* native to the British Isles, and *B. canariensis* native to the Canary Islands (Rasmont et al., 2008). The third taxon belongs to *B. terrestris* but its exact origin remains uncertain (hereafter *B. terrestris*). It is likely *B. terrestris ssp. dalmatinus* Dalla Torre or a hybrid between *B. terrestris ssp. dalmatinus* and *B. terrestris ssp. terrestris* (L.) (M. Pozo pers. comm.). These two taxa have a European continental distribution. Differences between subspecies within the *B. terrestris* aggregate include colouration and behavioural traits (Rasmont et al., 2008). The fourth taxon studied was *B. ignitus* Smith, which can be found in China, Korea and Japan (Shao et al., 2004).

We used five colonies from each the four taxa (six for *B. audax*). When the colonies were received (3 July 2018), each colony consisted of a single queen and approximately 20-40 workers. An additional colony of *B. audax* was added to the experiment on August 2018. Upon receipt, the colonies were placed under enclosed laboratory conditions at room temperature (20-23°C) and fed with *ad libitum* sugar solution from a plastic container placed under the colony (*Biogluc*, Biobest). Additionally, we provided each colony with 2 g of ground honeybee-collected pollen every week throughout the experiment. Bee experiments were conducted with approval from the Ethics Committee from the University of Stirling.

### Plant material

We studied two closely related species of buzz-pollinated plants in the genus *Solanum* (Solanaceae). *Solanum* flowers are nectarless and offer pollen as the main reward to attract pollinators. Pollen is contained within non-dehiscent poricidal anthers (Harris, 1905), which are arranged in the centre of the flower, sometimes forming a cone. The most efficient way to release pollen is through vibrations applied to the anthers (Bowers, 1975). *Solanum* is a classic system for the study of buzz-pollination (Buchmann et al., 1977; De Luca and Vallejo-Marín, 2013). Here we studied two annual species in the monophyletic clade *Solanum* Sect *Androceras* (Whalen, 1979): *S. rostratum* Dunal and *Solanum citrullifolium* (A.Braun) Nieuwl. These two species depart form the classical *Solanum*-type flower by having heterantherous flowers, in which the anthers are differentiated into two functional types (Vallejo-Marin et al., 2014) (Fig. 1). One type, the four feeding anthers, is yellow and placed at the centre of the flower, and the second type, the single pollinating anther, is usually differently coloured (yellow to reddish brown or light violet in the species studied here), and displaced to either the right- or left-hand side of the centre of the flower (Vallejo-Marin et al., 2014). At anthesis, the flowers are orientated with the anthers parallel to the ground. Bees usually manipulate the feeding anthers (Fig. 1), and only rarely directly manipulate the pollinating anther (Vallejo-Marin et al., 2009). Plants were grown from seed at the University of Stirling research glasshouses following the protocol described in Vallejo-Marín et al. (2014). Seeds from *S. rostratum* were collected in the field from open-pollinated fruits from three separate individuals (accessions 10s34, 10s36, and 10s39) in Puerto el Aire, Veracruz, Mexico (18.74°N, 97.53°W). Seeds from *S. citrullifolium* were obtained from self-fertilised fruits of three individuals (accession 199) grown from seeds originally obtained from the Radboud University Solanaceae collection (accession 894750197). Fresh flowers were transported from the glasshouse to the laboratory in open containers by cutting entire inflorescences and placing the stems in water-soaked Ideal Floral Foam (OASIS, Washington, UK). Inflorescences were then kept in wet floral foam and stored at room temperature until used for experiments.

### Bumblebee colony conditioning

The bumblebee colony conditioning was done in a 122cm × 100cm × 37cm wooden flight arena with a Perspex lid. The arena was divided in half with a wooden panel to allow up to two colonies to be connected at the same time. The arena was illuminated with a LED light panel (59.5cm × 59.5cm, 48W Daylight; Opus Lighting Technology, Birmingham, UK) on each side. To encourage bees to forage during the experiment, a pollen and a nectar feeder (Russell and Papaj, 2016) were placed in the centre of the flight arena. A single colony was attached to each side of the arena, and bees were allowed to enter the arena and return to the colony freely for at least one day. Following this, all colonies were exposed to a bouquet of *Solanum citrullifolium* and *Solanum rostratum* placed in the flight arena. After that period, we considered the colony “conditioned” and used it in buzz pollination trials.

### Buzz pollination trials

We conducted the buzz pollination trials in a 60cm × 60cm × 60cm “Russell” flight arena made of wooden panels and a UV-transparent Perspex lid, illuminated with a LED light panel on the top(A. Russell *pers. comm.*). The flight arena had two openings in the front side of the box covered with clear plastic to allow direct observations into the arena. A single bumblebee colony was connected at any one time to the flight arena using plastic tubes with sliding plastic/metal doors to control bee access to the arena. In each buzz pollination trial, a single bee was allowed to enter the flight arena and forage for a maximum of 10 min. If the bee did not visit the flower attached to the accelerometer (see below) during that time, we returned it to the colony and allowed the next bee to enter the arena. Bees that vibrated any flower were allowed to continue visiting the experimental array for up to 15 minutes to enable data collection, and subsequently captured and marked on the thorax with an individual ID using coloured pens (POSCA, Mitsubishi, Milton Keynes, UK). Marked bees were returned to the colony with their pollen baskets intact. Between trials, we replaced any flower that had been visited more than 3 times with fresh flowers.

In each buzz pollination trial, we placed an inflorescence with 1-3 flowers within the flight arena and recorded floral vibrations on a single focal flower using a calibrated miniature, lightweight (0.8 g) uniaxial accelerometer (Model 352A24, PCB Piezotronics Inc.). The accelerometer was attached to the focal flower through a small (5 × 0.35 mm) metallic pin glued with Loctite 454 to the accelerometer. The pin was then inserted into the base of the flower (receptacle) at a 90° angle to the longest axis of the stamens (Figure S1). Although the vibrations produced by the bee and transmitted to the flower likely result in movement of the flower in three dimensions (Gibson and Cocroft, 2018), our accelerometer only records vibrations along a single axis. We chose to measure vibrations along this axis, as theoretical models of buzz pollination suggest that pollen ejection is a function of the vibration magnitude along an axis perpendicular to the longest axis of the stamen (Buchmann and Hurley, 1978; Vallejo-Marin, in review). The accelerometer was connected to a battery-powered signal conditioner (480C02, PCB Piezotronics), and the signal was digitized using a digital storage oscilloscope (TBS1032B, Tektronix UK Limited) at a sampling rate of 2,500-25,000Hz (mean = 6,745Hz, median = 5,000Hz) and stored as a text file.

### Estimating vibrational properties

#### Accelerometer

Accelerometer recordings were imported to *R* v. 3.5.1 (R Development Core Team, 2018) and stored as time series objects. We selected from 1 to 4 recordings per bee and foraging bout. Sampling frequency and probe attenuation were extracted from each accelerometer file. The voltage readings of the accelerometer recorded in the oscilloscope were transformed to acceleration units (m/s^2^) using the product’s calibration (10.2 mV/ms^-2^). Time series were centred at zero and analysed using *seewave* (Sueur, 2018). In order to minimize low-frequency noise, we used a high-pass filter of 80 Hz (which is well below the fundamental frequency of bumblebee floral vibrations, De Luca and Vallejo-Marín, 2013) with a 512 Hamming Fourier Transform window using the function *fir*. We used the *timer* function to identify individual vibrations not broken by silence. A single vibration from each accelerometer file was randomly selected for extracting frequency and acceleration information (Fig. 2a). We calculated root mean squared (RMS) acceleration from the time-series data using the *rms* function (Fig. 2b). To calculate peak acceleration, we first obtained the amplitude envelope of the time wave (absolute amplitude) smoothed it with a window size of two and an overlap of 75%, and then obtained the maximum observed acceleration (Sueur, 2018) (Fig. 2c). This approach should provide a conservative estimate of peak acceleration that is more robust to voltage spikes in the time wave. Fundamental frequency was calculated over the duration of the vibration using the maximum possible window length (128—2048 samples, median = 512) within the limit imposed by the total number of samples of each vibration (window length ≤ number of samples). Larger window lengths improve frequency resolution (Δ_*f*_ = sampling frequency / window length) (Sueur, 2018; Vallejo-Marin, in review). We estimated the fundamental frequency using the function *fund* with a maximum of 500Hz (De Luca and Vallejo-Marín, 2013) and a window overlap of 75%. When the recording length allowed for multiple estimates of fundamental frequency for the same vibration, we calculated the mean and used this for analysis.

#### Coupling factor

The transmission of mechanical vibrations from the bee to the flower are likely influenced by the structural and material properties of the flower itself (Vallejo-Marin, in review). The vibrations we measured at the base of the flower are, therefore, potentially different from the vibrations experienced at the anthers. Moreover, interspecific differences in floral architecture make likely that the transmission of vibrations from the bee to the floral organs varies between plant species (Vallejo-Marin, in review). King (1993) suggests calculating a “coupling factor” to quantify the change in magnitude of the vibrations produced at the anther level and those measured away from the anthers. Following King (1993), we used a Bruel & Kjaer 4294 Calibration Exciter (Bruel & Kjaer, Naerum, Denmark) to produce vibrations of known frequency and acceleration (159.15 Hz ± 0.02%, 10 m/s^2^ ± 3% RMS acceleration). We applied the calibration exciter’s vibrations to the flower by firmly contacting the feeding anthers arranged in their natural position (Figure S1b). We recorded the vibrations experienced by the accelerometer attached to the flower’s calyx, and calculated the coupling factor, i.e., the ratio of RMS acceleration between the anther and calyx vibrations (coupling factor = RMS_callibration exciter_/RMS_accelerometer_). Values of the coupling factor greater than one indicate that the flower is reducing the amplitude of vibrations. RMS acceleration values were calculated as described in the ‘Measuring vibrational properties’ section. We estimated the coupling factor for each plant species using 30 flowers of *S. rostratum* (from 5 individuals and 3 accessions) and 35 flowers of *S. citrullifolium* (from 3 individuals and 3 accessions). We analysed an average of 11 recordings per flower (N = 365 for *S. citrullifolium* and N = 370 for *S. rostratum*). RMS acceleration was calculated as above.

#### Peak displacement

We estimated peak displacement using fundamental frequency and peak acceleration, assuming simple sinusoidal motion, using the equation *PD = PA*/(2π*F*)^2^, where *PD* is peak displacement, *PA* is peak acceleration amplitude, and *F* is fundamental frequency (c.f. King and Buchmann, 1996). To estimate peak displacement incorporating the coupling factor, we used the value of PA multiplied by the species-specific coupling factor calculated as above.

#### Flower mass

As the mass of the flower could influence the magnitude of the vibrations transmitted from the body of the bee to the floral receptacle, we estimated flower mass of each flower used in the buzz pollination trials. In order to estimate flower mass before visitation (and hence before pollen removal) we used an indirect approach. We randomly sampled 100 flowers of each plant species (including the pedicel) and used them to estimate a species-specific correlation between flower morphological measurements and dry biomass (mg). To do this, we measured maximum corolla length, the length of the feeding anthers and the pollinating anther length. These three morphological measurements were summarised using a principal component analysis, and the first principal component (PC1) was used as an overall estimate of flower size. Then, each flower was dried at 60°C for 36 hours and weighted with an analytical balance with a 0.01mg resolution (Analytical Plus AP250D, Ohaus, Greifensee, Switzerland). We then calculated a correlation between PC1 and flower mass (mg) and used this association to calculate the dry biomass of experimental flowers from morphological measurements.

#### Bee size

As an estimate of bee size, we used intertegular distance (ITD), which has been shown to be a good approximation of overall bee size (Cane, 1987), and is regularly used in buzz-pollination studies (De Luca et al., submitted). At the end of the behavioural trials (approximately end of August), we euthanised bees by placing the entire colony in a -80°C freezer for 48h. All marked bees and an additional random sample of workers, all identifiable males, and queens were measured (100 bees per species, 20 per colony). Bees (workers only) were measured with digital callipers (Mitutoyo, Andover, UK) to the nearest 0.1 mm. In total, we measured 557 worker bees from 21 colonies of four taxa.

### Statistical analysis

To compare differences in size (intertegular distance, ITD) between workers of different bee species, we used a linear mixed-effects model with ITD as a response variable, species as an explanatory variable (fixed effect) and colony as a random effect (21 colonies, N = 557 bees). We used a Tukey test for pairwise comparisons between bee taxa. To compare the size of bees that were observed producing floral vibrations (buzzers) vs. a random sample of workers from each colony (all bees), we selected colonies that included both categories of bees (8 colonies, N = 282 bees). In this analysis, ITD was used as a response variable, bee species and bee type (buzzer vs. all bees) as fixed effects, and colony as a random effect. The interaction between species and bee type was excluded during preliminary analyses as it was not significant. All bees included worker bees that left the colony and did not buzz, bees that never left the colony, and may have included some bees that buzzed but were not observed and marked.

To test for differences in the vibrational properties between bumblebee species and between flowers of the two *Solanum* species, we also used linear mixed-effects models. The response variables were either RMS acceleration, peak acceleration, fundamental frequency or peak displacement. Bee size (ITD), flower mass, bumblebee species and plant species were included as fixed effects. As we had several vibrational measures for most bumblebee individuals, we used bumblebee individual as a random effect, which accounts for the statistical associations among vibrations produced by the same individual. All analyses were done in *R* v.3.5.1 (R Development Core Team, 2018) using packages *lme4* (Bates et al., 2014) for parameter estimation and *lmerTest* (Kuznetsova et al., 2014) for assessing statistical significance.

## Results

We found that size (ITD) of individual worker bees was significantly different among species (*P*< 0.01; Figures S2, S3). Specifically, *Bombus audax* was smaller than the other three taxa, which did not differ significantly in size from each other (Figure S2). A comparison of the size of bees observed vibrating *Solanum* flowers against all worker bees sampled within each colony (using colony as a random effect and bee species as a fixed effect), showed that buzz-pollinating bees were on average larger (coefficient for all workers = -0.286, *P* < 0.001) (Figure 3).

**Figure 3.**
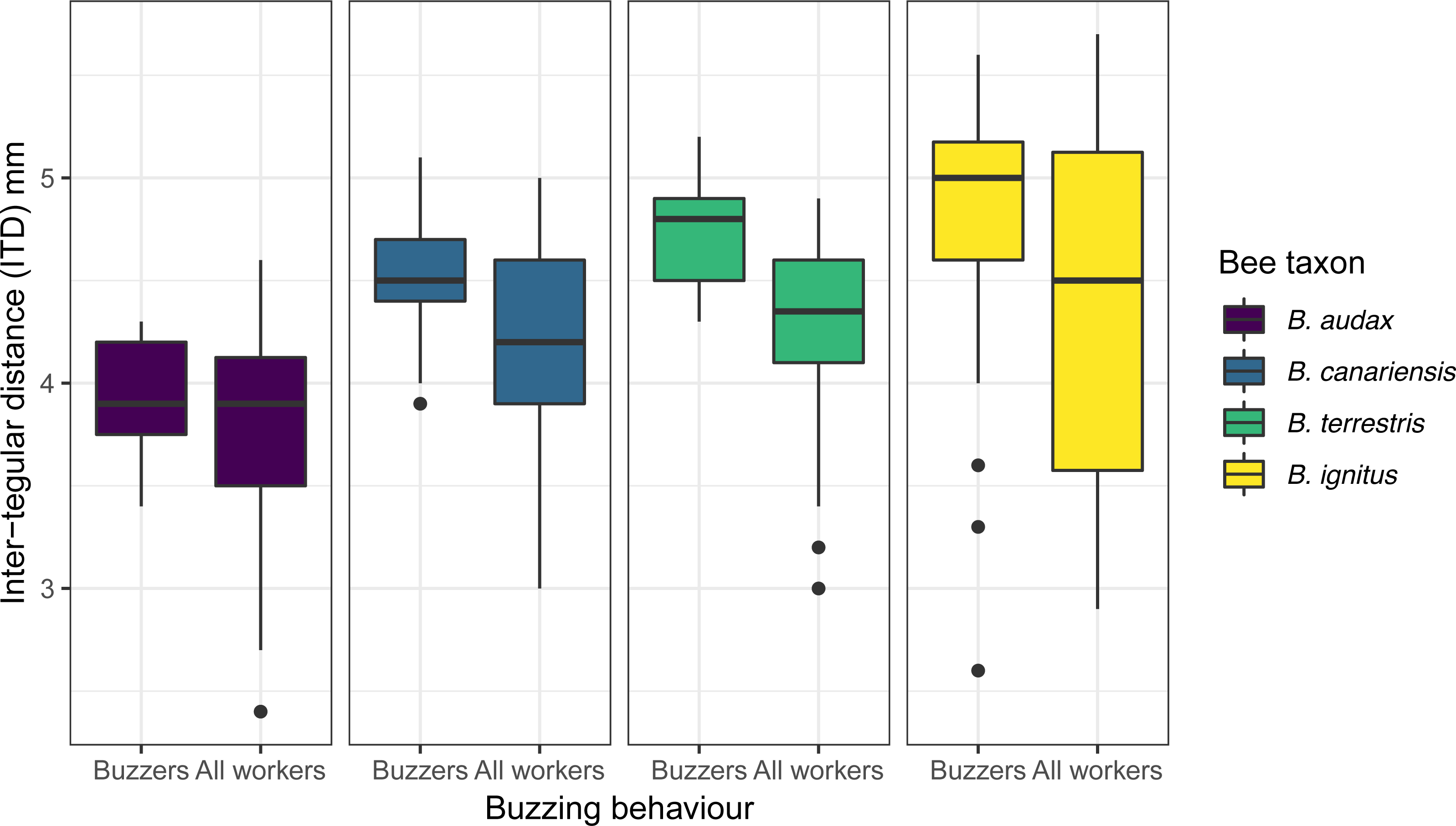
Size (intertegular distance, mm) of bees observed buzzing during the experiment (buzzer) vs. a random sample of workers in the colony (all workers). This analysis included only colonies in which both buzzer and all workers were observed and measured during the experiment (8 colonies, two per taxon) (N = 282 bees).

We obtained a total of 230 recordings of floral vibrations from 79 different individual bumblebees from 8 colonies of four different taxa (Table 1). All colonies and bee taxa were recorded on *S. rostratum*, while three of those colonies (two *Bombus audax* and one *B. canariensis* colony) were recorded both in *S. rostratum* and on *S. citrullifolium* (102 recordings on *S. rostratum* and 82 on *S. citrullifolium*). The analysis of calibrated vibrations applied to the anthers and measured with the accelerometer at the base of the flower, showed that the two studied plant species differed in the damping of vibrations transmitted through the flower. The coupling factor for *S. rostratum* was 8.84 ± 0.23 (mean ± SE) and for *S. citrullifolium* was 12.21 ± 0.29, indicating that flower of *S. citrullifolium* dampen floral vibrations more strongly than those of its congener. The observed differences in the coupling factor of each plant species were statistically significant as assessed with a linear model with species as explanatory variable (species coefficient (*S. rostratum*) = -3.367, *P*< 0.001).

**Table 1.**
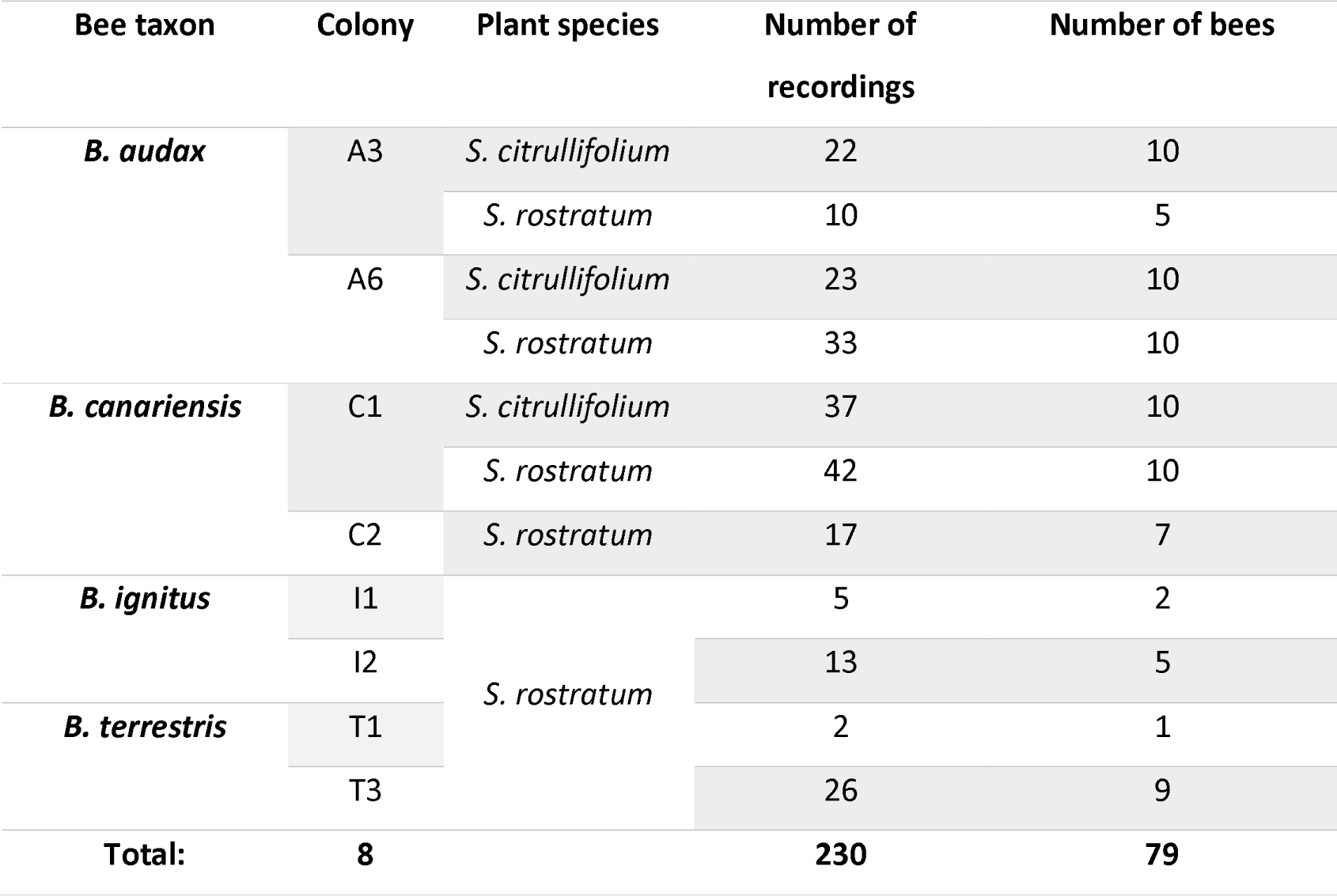
Sample size for the experiments measuring floral vibrations. Number of recordings represents the number of accelerometer recordings that were used in the final analysis. The number of bees indicate the number of individuals used in each plant species/bee colony combination. A few of the same individuals were recorded in both plant species, and thus the total number of different bees across the whole experiment was 72.

## Differences between bumblebee taxa on *Solanum rostratum*

We found significant differences among bumblebee species in peak acceleration, RMS acceleration (with and without coupling factor) and fundamental frequency (Table 2, Figure 4). *B. audax* and *B. terrestris* had floral vibrations with the highest fundamental frequency, followed by *B. canariensis* and *B. ignitus* (Figure 4a). *B. audax* had a RMS acceleration higher than all other taxa, as well as one of the highest peak accelerations (with *B.* ignitus) (Figs. 4b and 4c). The other three taxa achieved similar levels of both RMS and peak acceleration. Intertegular distance and flower biomass were not significantly associated with either measurement of acceleration or with frequency (Table 2). Peak displacement calculated using fundamental frequency and peak acceleration was significantly different among bee species. Peak displacement was higher for *B. ignitus* and *B. audax*, and lower for *B. canariensis* and *B. terrestris* (Table 2; Fig. 4b).

**Table 2.** Effect of bee taxon, bee size (intertegular distance) and floral mass on the mechanical properties of floral vibrations produced by four bee taxa visiting the same plant species (*Solanum rostratum*). Floral vibrations produced by four bumblebee taxa on flowers of buzz-pollinated *Solanum rostratum* were recorded with an accelerometer attached to the base of the flower. Peak acceleration, root mean square (RMS) acceleration, and fundamental frequency were calculated for a subset of randomly selected floral vibrations. Parameter estimates and standard errors for individual coefficients were obtained from a linear mixed-effects model with bee individual as a random effect. *P*-values were calculated for each explanatory variable using a Type III analysis of variance with Sattertwhaite’s method.

|  | Peak acceleration (m/s<sup>2</sup>) |  |  | RMS acceleration (m/s<sup>2</sup>) |  |  | RMS acceleration, with flower coupling factor (m/s<sup>2</sup>) |  |  |
| --- | --- | --- | --- | --- | --- | --- | --- | --- | --- |
| Parameters: | Estimate | SE | P-value | Estimate | SE | P-value | Estimate | SE | P-value |
| Intercept (*B. audax*) | 1.908 | 7.689 |  | 1.411 | 4.953 |  | 12.477 | 43.788 |  |
| Intertegular distance (mm) | 1.032 | 1.597 | 0.521 | 0.938 | 1.079 | 0.389 | 8.294 | 9.538 | 0.389 |
| Floral dry mass (mg) | 0.392 | 0.241 | 0.106 | 0.127 | 0.139 | 0.361 | 1.125 | 1.227 | 0.361 |
| Bee taxon |  |  | 0.015 |  |  | 0.003 |  |  | 0.003 |
| *B. canariensis* | -5.291 | 1.556 |  | -4.059 | 1.064 |  | -35.886 | 9.409 |  |
| *B. terrestris* | -5.348 | 1.995 |  | -4.506 | 1.355 |  | -39.831 | 11.976 |  |
| *B. ignitus* | -4.268 | 2.016 |  | -4.140 | 1.382 |  | -36.594 | 12.219 |  |

| Fundamental frequency (Hz) |  |  | Peak displacement (μm) |  |  |
| --- | --- | --- | --- | --- | --- |
| Estimate | SE | P-value | Estimate | SE | P-value |
| 421.686 | 44.406 |  | -0.969 | 1.729 |  |
| -1.768 | 8.818 | 0.842 | 0.390 | 0.345 | 0.265 |
| -2.696 | 1.482 | 0.071 | 0.109 | 0.057 | 0.058 |
|  |  | **<0.0001** |  |  | **<0.001** |
| -40.565 | 8.460 |  | -0.707 | 0.332 |  |
| -3.418 | 10.992 |  | -1.288 | 0.430 |  |
| -93.305 | 10.990 |  | 0.622 | 0.431 |  |

**Figure 4.**
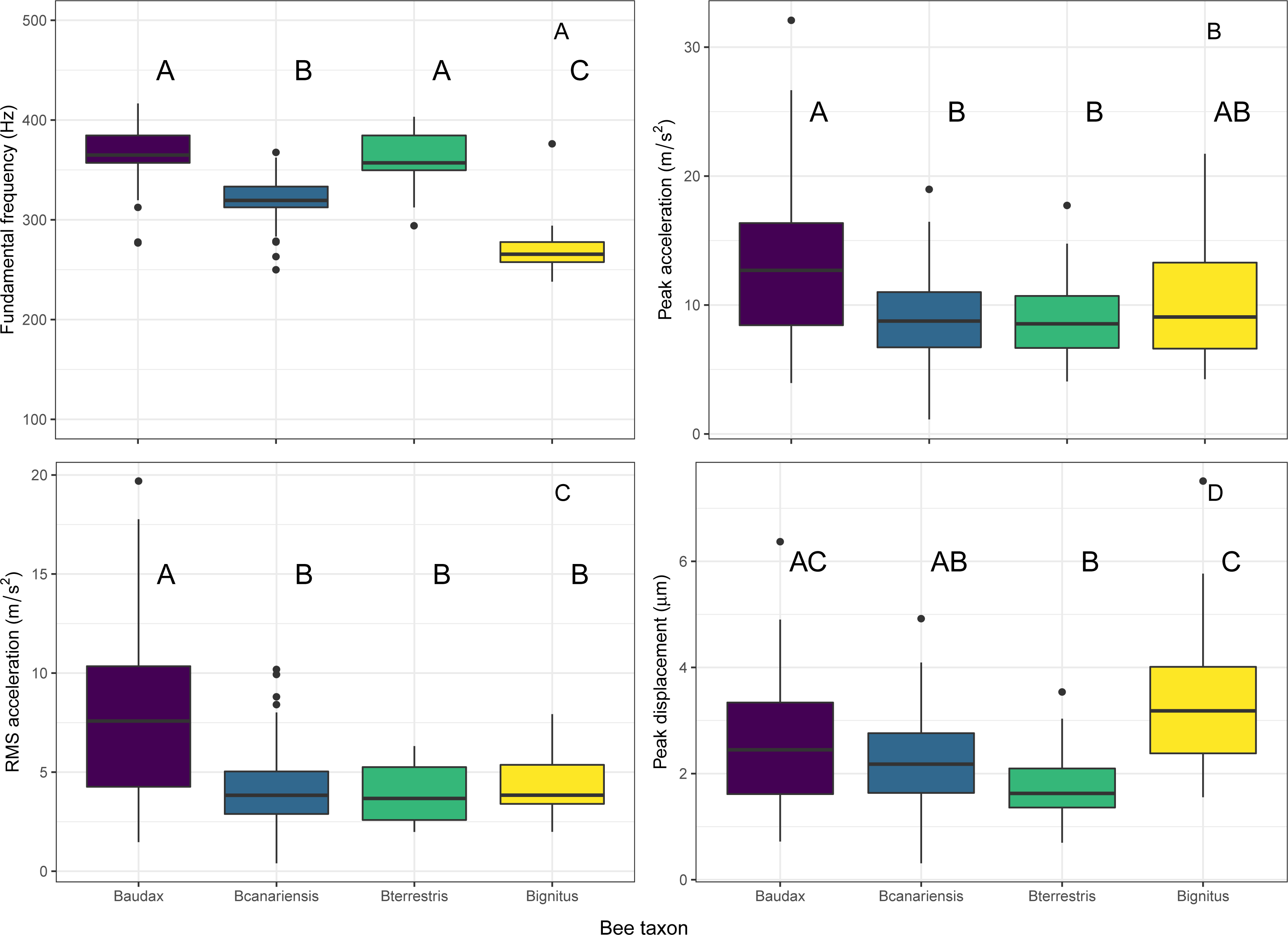
Comparison of the mechanical properties of floral vibrations from four taxa of bumblebees on flowers of buzz-pollinated *Solanum rostratum* (Solanaceae). Vibrations were recorded using an accelerometer attached to calyx of the flower using a metallic pin. **(A)** Fundamental frequency. **(B)** Peak acceleration. **(C)** Root Mean Square (RMS) acceleration. **(D)** Peak displacement calculated with RMS acceleration. Within each panel, different letters indicate statistically different mean values assessed by a Tukey test of pairwise comparisons with a confidence level of 95%. N = 148 floral vibrations from 49 bees from 8 colonies of four taxa. Baudax *= Bombus terrestris ssp. audax,* Bcanariensis *= B. terrestris ssp. canariensis, Bignitus = B. ignitus,* Bterrestris *= B. terrestris.*

## Differences between bee taxa visiting two plant species

We found a statistically significant effect of both bee taxon (*B. audax* vs. *B. canariensis*) and plant species (*S. rostratum* vs. *B. citrullifolium*) on peak and RMS acceleration of floral vibrations (Table 3, Fig. 5). In contrast, when we multiplied RMS acceleration by the coupling factor calculated for each *Solanum* species, bee species still contributed significantly to explaining acceleration values but the effect of plant species on acceleration disappeared. Fundamental frequency was also significantly different between bee taxa, but not between plant species (Table 3, Fig. 5). Neither bee size not flower biomass were significantly associated with the acceleration of floral vibrations (Table 3). Unexpectedly, bee size (ITD) was positively associated with the fundamental frequency of floral vibrations after statistically accounting for differences in size between the two bumblebee taxa (Table 3). Peak displacement was significantly different between plant species but not between *B. audax* and *B. terrestris*. However, the difference in peak displacement between plants disappeared when accounting for the plant’s coupling factor (Table 3, Fig. 6).

**Table 3.**
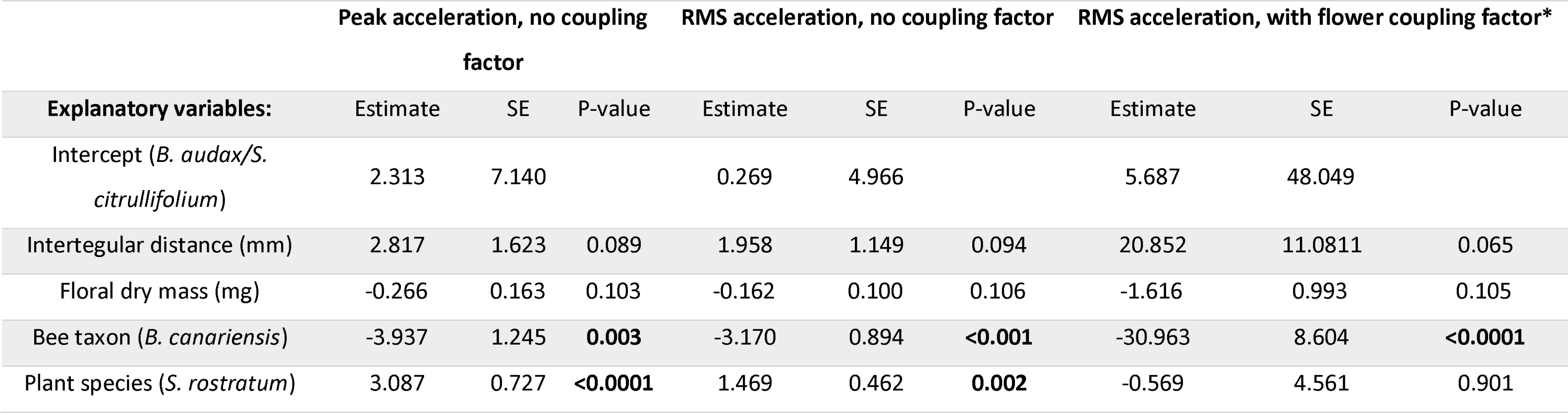
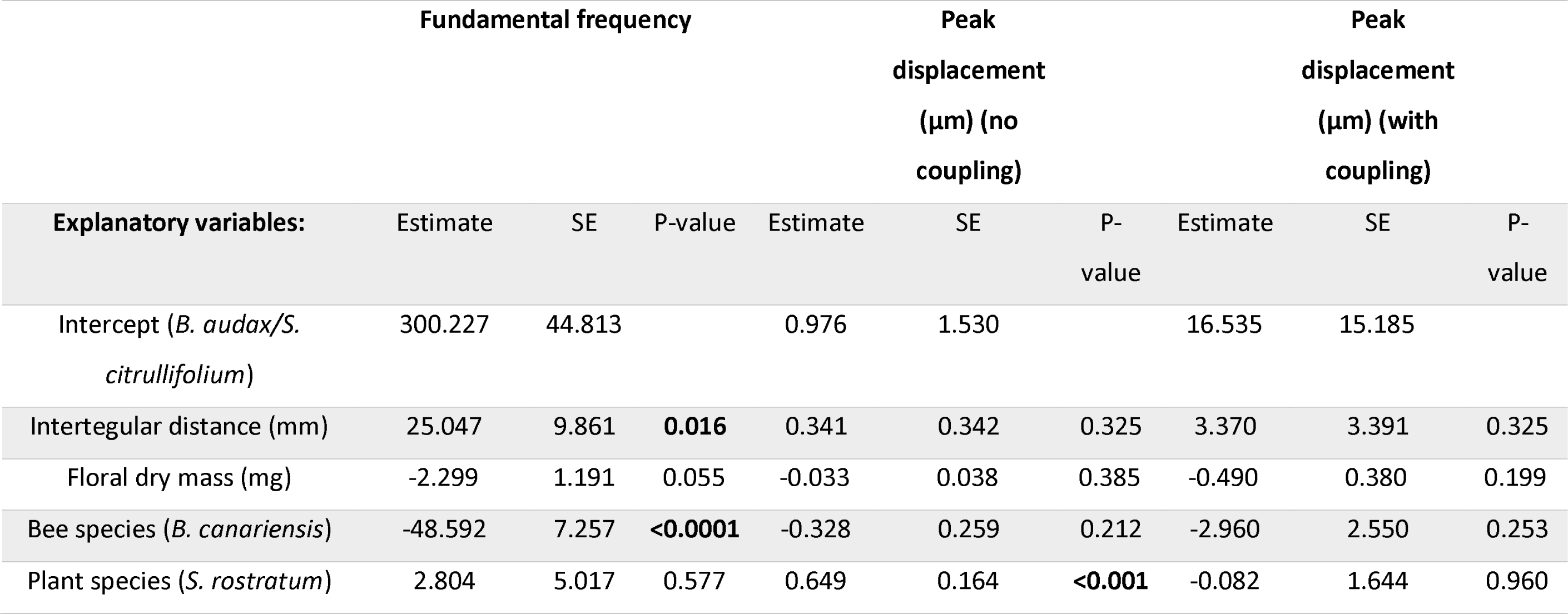
Effect of bee taxon, bee size (intertegular distance), plant species and floral mass on the mechanical properties of floral vibrations during buzz pollination of two plant species (*S. rostratum* and *S. citrullifolium*) visited by two bee taxa (*B. audax* and *B. canariensis*). Floral vibrations produced by four bumblebee taxa on flowers of buzz-pollinated Solanum rostratum were recorded with an accelerometer attached to the base of the flower. Peak acceleration, root mean square (RMS) acceleration, and fundamental frequency were calculated for a subset of randomly selected floral vibrations. Parameter estimates and standard errors for individual coefficients were obtained from a linear mixed-effects model with bee individual as a random effect. *P*-values were calculated for each explanatory variable using a Type III analysis of variance with Sattertwhaite’s method.*In this analysis, the response variable is peak acceleration multiplied by a species-specific flower coupling factor. Flower coupling factor: *S. rostratum* = 8.56; *S. citrullifolium* = 11.30.

|  | Peak acceleration, no coupling factor |  |  | RMS acceleration, no coupling factor |  |  | RMS acceleration, with flower coupling factor* |  |  |
| --- | --- | --- | --- | --- | --- | --- | --- | --- | --- |
| Explanatory variables: | Estimate | SE | P-value | Estimate | SE | P-value | Estimate | SE | P-value |
| Intercept (*B. audax*/*S. citrullifolium*) | 2.313 | 7.140 |  | 0.269 | 4.966 |  | 5.687 | 48.049 |  |
| Intertegular distance (mm) | 2.817 | 1.623 | 0.089 | 1.958 | 1.149 | 0.094 | 20.852 | 11.0811 | 0.065 |
| Floral dry mass (mg) | -0.266 | 0.163 | 0.103 | -0.162 | 0.100 | 0.106 | -1.616 | 0.993 | 0.105 |
| Bee taxon (*B. canariensis*) | -3.937 | 1.245 | **0.003** | -3.170 | 0.894 | **<0.001** | -30.963 | 8.604 | **<0.0001** |
| Plant species (*S. rostratum*) | 3.087 | 0.727 | **<0.0001** | 1.469 | 0.462 | **0.002** | -0.569 | 4.561 | 0.901 |

| Explanatory variables: | Fundamental frequency |  |  | Peak displacement (μm) (no coupling) |  |  | Peak displacement (μm) (with coupling) |  |  |
| --- | --- | --- | --- | --- | --- | --- | --- | --- | --- |
|  | Estimate | SE | P-value | Estimate | SE | P-value | Estimate | SE | P-value |
| Intercept (*B. audax*/*S. citrullifolium*) | 300.227 | 44.813 |  | 0.976 | 1.530 |  | 16.535 | 15.185 |  |
| Intertegular distance (mm) | 25.047 | 9.861 | **0.016** | 0.341 | 0.342 | 0.325 | 3.370 | 3.391 | 0.325 |
| Floral dry mass (mg) | -2.299 | 1.191 | 0.055 | -0.033 | 0.038 | 0.385 | -0.490 | 0.380 | 0.199 |
| Bee species (*B. canariensis*) | -48.592 | 7.257 | **<0.0001** | -0.328 | 0.259 | 0.212 | -2.960 | 2.550 | 0.253 |
| Plant species (*S. rostratum*) | 2.804 | 5.017 | 0.577 | 0.649 | 0.164 | **<0.001** | -0.082 | 1.644 | 0.960 |

**Figure 5.**
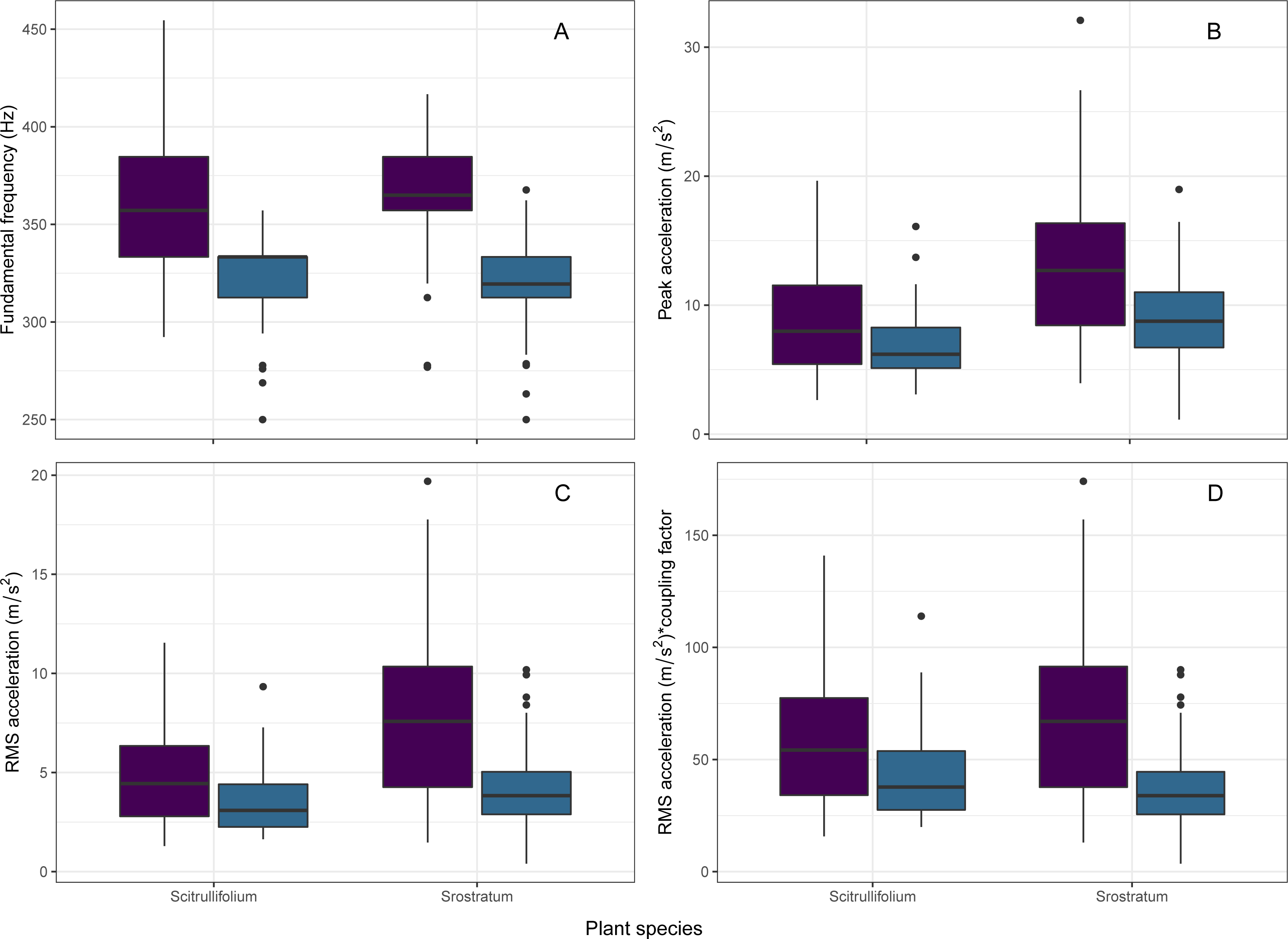
Comparison of the mechanical properties of floral vibrations from two bumblebee taxa (*Bombus terrestris ssp. audax* and *B. terrestris ssp. canariensis*) on flowers of two buzz-pollinated plants (*Solanum citrullifolium* and *S. rostratum*). Vibrations were recorded using an accelerometer attached to calyx of the flower using a metallic pin. **(A)** Fundamental frequency. **(B)** Peak acceleration. **(C)** Root Mean Square (RMS) acceleration. **(D)** RMS acceleration multiplied by the species-specific flower coupling factor (see Methods). N = 184 floral vibrations from 53 bees from four colonies of two taxa. Purple = *Bombus terrestris ssp. audax*, blue = *B. terrestris ssp. Canariensis*

**Figure 6.**
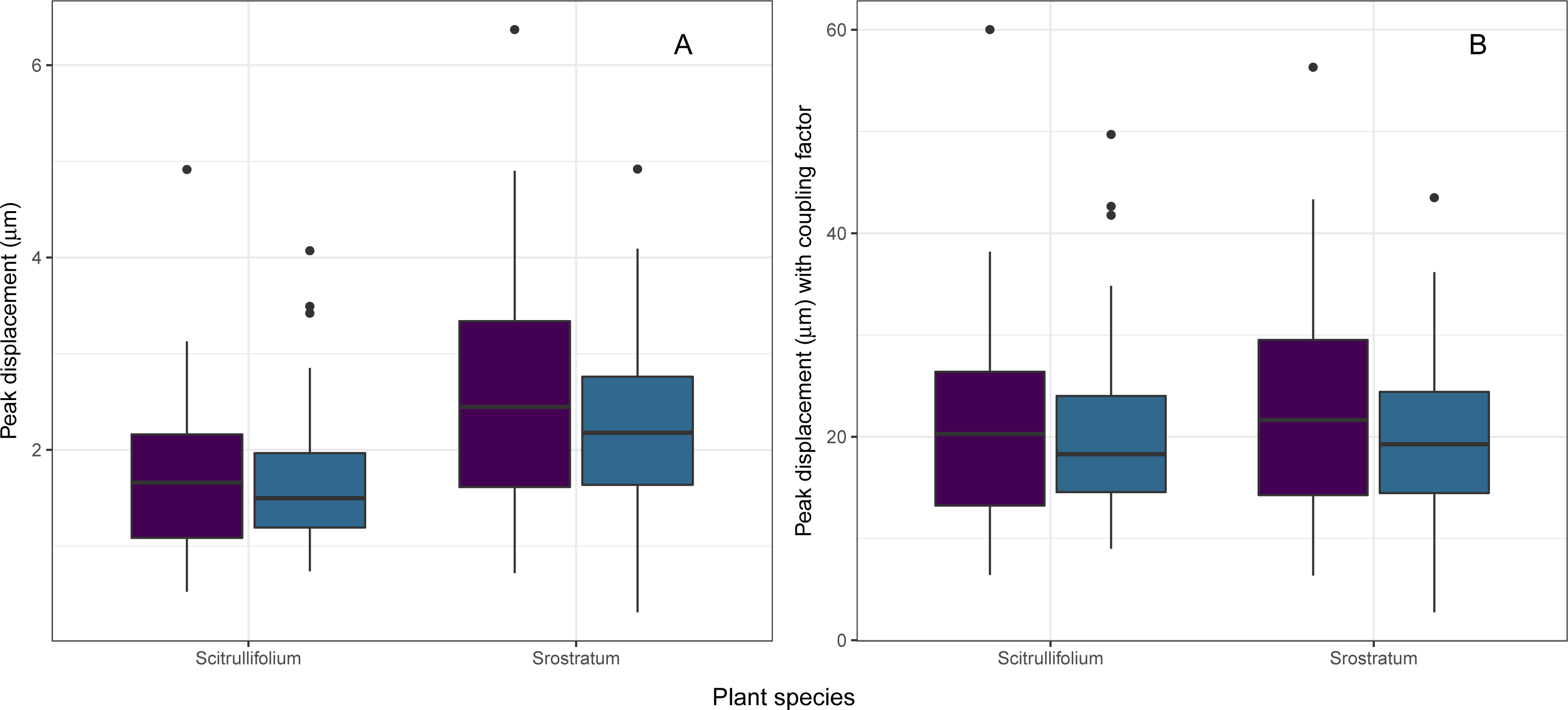
Estimated peak displacement (µm) of floral vibrations produced by two bumblebee taxa (*B. audax*, purple; *B. canariensis*, blue) while visiting flowers of two plant species (*S. citrullifolium* and *S. rostratum*). **(A)** Estimated peak displacement at the point of measurement, i.e., the base of the flower. **(B)** Estimated peak displacement at the anthers, obtained by multiplying peak acceleration by the plant species-specific coupling factor. N = 184 floral vibrations from 53 bees from four colonies of two taxa.

## Discussion

Our study represents one of the few available investigations on the mechanical properties of floral-borne vibrations produced by bees during buzz pollination, and one of the first to systematically compare substrate-borne vibrations of multiple bee taxa on the same flower, and of the same bee taxon on different plant species. We found that floral vibrations on the same plant species (*S. rostratum*) differ between bee taxa in both fundamental frequency and acceleration amplitude (both RMS acceleration and peak acceleration). Interestingly, although neither individual bee size nor floral biomass explained variation in floral vibration properties, *audax*, the bee taxon with the smallest average size, produced floral vibrations that combined both relatively high frequency and acceleration. When measurements of frequency and acceleration were used to calculate peak displacement (the maximum distance travelled by the oscillating structure from its resting position), we found statistically significant differences between bee taxa, with both *B. audax*, and *B. ignitus* achieving larger displacements than *B. terrestris*. A comparison of floral vibrations produced by *B. audax* and *B. canariensis* visiting the same two *Solanum* plant species showed that acceleration (RMS and peak) and peak displacement measured at the base of the flower, but not frequency, are significantly different between plants. Comparison of the plant-specific coupling factors clearly indicated that flowers of even closely related species differ significantly in their capacity to dampen substrate-borne vibrations. We found that when these differences in transmission properties of flowers are taken into account, plant-specific differences in acceleration and displacement disappeared. Together, our results suggest that both *B. audax* and *B. canariensis* produce vibrations at the anther level that achieve similar acceleration amplitude and displacement irrespectively of the plant they visit. The main findings of our study can be summarised in three points: (1) Even closely related, and morphologically similar, bumblebee taxa differ in the mechanical properties of the vibrations transmitted to flowers during buzz pollination. (2) Floral type (i.e., plant species) significantly influences the acceleration amplitude of vibrations transmitted over floral structures. (3) *B. audax* and *B. canariensis*, two closely related subspecies within the *B. terrestris* species aggregate, produce approximately similar acceleration and displacement while vibrating the anthers of two different buzz-pollinated plants.

## Differences in floral vibrations between closely related bees

Previous studies have shown that different species of bees produce vibrations with different properties when sonicating flowers of the same plant species (e.g., Burkart et al., 2011; Corbet and Huang, 2014; King and Buchmann, 2003). Most of these previous studies have focused on the properties of the acoustic (airborne) component of floral vibrations, particularly frequency, in part due to the ease and accessibility of recording acoustic vibrations (De Luca et al., 2018). Differences in the frequency of floral vibrations might be expected given size variation among buzz pollinating bees. It is well established that wingbeat frequency in insects is correlated with morphological characteristics including wing length and body mass (Byrne et al., 1988), and similarly, body mass is associated with frequency in insects communicating using airborne sound (Cocroft and De Luca, 2006). However, floral vibrations seem to be less strongly associated with specific morphological traits in bees, such as body size (Burkart et al., 2011; De Luca et al., submitted; Rosi-Denadai et al., 2018).

Here we found that although different bee species produced floral vibrations with different properties, body size (ITD) did not consistently explain variation in either the frequency or amplitude of substrate-borne vibrations. In the first experiment (multiple bee species in the same plant), we found no association between bee size and frequency or amplitude. In the second experiment comparing vibrations of two bee species in *S. rostratum* and *S. citrullifolium*, we found that bee size (ITD) was positively correlated with frequency but not with amplitude. A positive association between body size and frequency is contrary to the negative associated usually found between body size and wingbeat frequency (De Luca et al., submitted). The lack of a consistent relationship between body size and frequency is paralleled by a previous study of buzz-pollinating *B. impatiens* (Switzer and Combes, 2017). The authors of this study found that higher (acoustic) frequency during floral vibrations was positively associated with bee’s body mass, but negatively associated with ITD. In a study of the characteristics of substrate-borne vibrations used in insect communication, Cocroft and De Luca (2006) found a negative relationship between body size and signal frequency across 51 species in the Family Membracidae (Hemiptera), but no correlation when analysing within-population variation in two species of treehoppers. Importantly, they also found that the association between body size and signal frequency across species of different orders varied with the function of those signals (e.g., alarm or mating). Our results can be interpreted as to say that, after accounting for the effect of bee species, size variation within species does not correlate with the amplitude of floral vibrations although it may be positively associated with frequency in some plants. This has some interesting implications for buzz pollination, as it suggests that within the range of size variation seen among buzz-pollinating workers bees of given species, different-sized workers are capable of producing similar floral vibrations. This could be achieved either passively (floral vibrations do not depend strongly on bee traits related to size) or actively (bees may be able to adjust their vibrations to achieve a given outcome). To the extent to which the properties of floral vibrations affect pollen removal (De Luca et al., 2013), this should also influence the amount of pollen obtained by workers. More research is needed to establish the morphological, physiological and behavioural determinants of floral vibrations, and the question as to why even closely-related species differ in their floral vibrations remains unanswered.

## Do bees adjust their vibrations depending on plant species?

Previous studies indicate that the characteristics of floral vibrations of the same bee species on different plant species can vary (reviewed in Switzer and Combes, 2017). For example, in a greenhouse study of *B. impatiens* visiting three species of *Solanum* (*S. carolinense, S. dulcamara* and *S. lycopersicum*), Switzer and Combes (2017) showed that the same individual bee can produce floral vibrations of different frequency and duration on different plant species. We found that the acceleration amplitude, but not fundamental frequency, experienced at the base of the flower when buzz-pollinated by the same bee species varied depending on plant type. In contrast to previous studies measuring the frequency of airborne vibrations during buzz pollination, here we found that two species of *Bombus* produce substrate-borne vibrations of similar frequency while visiting two closely-related species of *Solanum*. In our study, the difference in floral vibrations among plants is restricted to changes in vibration amplitude, specifically acceleration amplitude.

The observed difference in vibration characteristics detected at the base of the flower of two plant species could have at least two explanations (Switzer and Combes, 2017). First, the bee may be actively adjusting its floral vibrations depending on plant species. Individual bees seem to be capable of modifying the frequency of their floral vibrations on a given plant species (Morgan et al., 2016), and it has been suggested that differences in floral characteristics including morphology and pollen availability may cause bees to change the vibrations produced by the same bee on different plants (Switzer and Combes, 2017). Second, differences in the characteristics of the substrate-borne vibrations measured here could be due to intrinsic differences in the mechanical properties of the flowers of the two species of *Solanum* studied. Our experimental design and analysis allowed us to calculate the mechanical changes to known vibrational signals as they travelled through the flower (the coupling factor). The analysis of floral vibrations after accounting for this plant species-specific differences indicate that a given bee species maintains similar types of vibrations regardless of the plant being visited. Thus, our study suggests that the differences in floral vibrations observed here are due to differences in the mechanical properties of flowers rather than to behavioural changes in the production of vibrations by bees. It remains a distinct possibility that the lack of amplitude differences in floral vibrations on different plants observed, result from the fact that bees are vibrating flowers are at, or close to, the maximum amplitude they can generate using their thoracic muscles.

In nature, the coupling between the bee and the rest of the flower is influenced by both the characteristics of the flower and by the way in which a bee holds onto floral parts while buzz-pollinating (King, 1993). Our experiment could only account for differences in the transmission of floral vibrations due to the mechanical properties of flowers, as the calibrated vibrometer used here cannot accommodate subtle variation in how a bee holds a flower during buzz pollination. More work is needed to determine how the way in which a bee manipulates a flower, e.g., how “tightly” does the bee holds the anthers or whether a bee “bites” the anthers (e.g., Russell et al., 2016), affects the transmission of vibrations. Different floral morphologies may cause changes in the way in which a bee manipulates and vibrates a flower, and therefore on the transmission of those vibrations. Both species of *Solanum* studied here have roughly similar floral morphologies and were manipulated in the same manner by bumblebees, but comparisons of floral species with more disparate morphologies may reveal if, and to what extent, changes in bee behaviour affect the transmission of floral vibrations, including vibration amplitude, and potentially pollen release.

## Effect of floral properties on substrate-borne floral vibrations

The field of animal communication using substrate-borne vibrations (biotremology) has long recognised the importance of considering plant characteristics on the functional ecology and evolution of vibrational communication (Cocroft and Rodríguez, 2005; Michelsen et al., 1982; Mortimer, 2017). In buzz pollination, the importance of plant traits in the transmission of vibrations is recognised (e.g., Buchmann and Hurley, 1978; King, 1993) but has rarely been explicitly incorporated into mechanistic, ecological or evolutionary studies.

Characteristics of the floral structures with which bees interact during buzz pollination, e.g., the androecium, should have an effect on the transmission of vibrational energy from the bee to the flower (King and Buchmann, 1996). Bees often vibrate only some of the anthers on a flower, but their vibrations can be transmitted to other floral parts, including other stamens not directly in contact with the bee, resulting in pollen release (MVM, per. obs.). Biotremology studies show that plants can cause significant distortion of insects’ vibration signals (Cocroft and Rodríguez, 2005; Michelsen et al., 1982). The characteristics of vegetative and flower tissues are therefore likely to cause alterations in the transmission of vibrations (King and Buchmann, 1996). Our results suggest that mass could not explain differences between flowers from two *Solanum* species when vibrating by the same *Bombus* species. However, we showed that *Solanum rostratum* reduced the amplitude of the vibrations transmitted through the flower less than *S. citrullifolium*. The transmission of vibrations through plant tissue depends not only on the distance from the source but also on the characteristics and vibration modes of the plant (Michelsen et al., 1982). Even fine structural differences, such as the presence of vascular tissue and vein size can affect the damping of vibrations (Michelsen et al., 1982). Thus, floral properties such as the morphology, geometry and material properties of the flower are expected to contribute to vibrational differences. For example, the stiffness of the filament holding the anther and whether the anther cones are loosely arranged or tightly packed into a cone (Glover et al., 2004), likely affect the transmission of vibrations away from its source (the bee). The vibrational properties of flowers particularly in an evolutionary context are not well understood. The convergence of disparate plant groups onto similar buzz-pollinated floral morphologies (e.g., the *Solanum*-type flower has evolved in more than 21 plant families; De Luca and Vallejo-Marín, 2013) provides an opportunity to investigate whether morphological similarity is associated with similar vibrational properties.

## Conclusions

Together with previous work, our results suggest that bees differ in their capacity to transmit vibrations to flowers. Because the characteristics of vibrations, in particular their amplitude, is associated with pollen release (De Luca et al., 2013; Vallejo-Marin, in review), visitation by different species of bees may affect patterns of pollen dispensing, and, ultimately, plant fitness (Harder and Barclay, 1994; Harder and Wilson, 1994). Therefore, from a plant’s perspective it may be selectively advantageous to favour visitation by specific visitors. Similarly, from a bee’s perspective the choice of a given plant species may determine the extent to which their floral vibrations translate into sufficient vibrational energy to elicit pollen release (Buchmann and Hurley, 1978). Particularly if bees cannot adjust their floral vibrations depending on plant species, selectively foraging in plants with low coupling factors, may allow bees to maximise pollen release per buzz effort. Moreover, bee traits that minimise the bee’s coupling factor (e.g., the location and manner in which a bee holds a flower) should also be favoured as mechanisms to transmit vibrations to the flowers more efficiently, potentially removing more pollen.

## Acknowledgements

We thank M. Pozo and Biobest for providing the bumblebee colonies for this experiment. A. Russell kindly shared his protocols and advice on flight arena construction and bee experiments in general. J. Gibson and P. De Luca provided key advice on the analysis and interpretation of substrate-borne vibrations and on buzz pollination. We thank J. Weir for help with flight arena construction. Members of the Vallejo-Marin Lab, in particular Lucy Nevard, provided help with bee maintenance. This study was partially funded by a SPARK grant (University of Stirling) to MVM, and an ERASMUS internshipto BAC.

## Data availability

Data will be deposited in Dryad and made publicly available upon publication.

